# Diminished ventral oligodendrocyte precursor generation results in the subsequent over-production of dorsal oligodendrocyte precursors of aberrant morphology and function

**DOI:** 10.1101/2020.03.09.983908

**Authors:** Lev Starikov, Andreas H. Kottmann

## Abstract

Oligodendrocyte precursor cells (OPCs) arise sequentially first from a ventral and then from a dorsal precursor domain at the end of neurogenesis during spinal cord development. Whether the sequential production of OPCs is of physiological significance has not been examined. Here we show that ablating Shh signaling from nascent ventricular zone derivatives and partially from the floor plate results in a severe diminishment of ventral derived OPCs but normal numbers of motor neurons in the postnatal spinal cord. In the absence of ventral vOPCs, dorsal dOPCs populate the entire spinal cord resulting in an increased OPC density in the ventral horns. These OPCs take on an altered morphology, do not participate in the removal of excitatory vGlut1 synapses from injured motor neurons, and exhibit morphological features similar to those found in the vicinity of motor neurons in the SOD1 mouse model of Amyotrophic Lateral Sclerosis (ALS). Our data indicates that vOPCs prevent dOPCs from invading ventral spinal cord laminae and suggests that vOPCs have a unique ability to communicate with injured motor neurons.

## Introduction

Oligodendrocyte precursor cells (OPCs) are a heterogeneous, proliferative, and highly mobile population of glia in the adult central nervous system [1–3]. OPCs tile the entire central nervous system (CNS) and give rise to myelinating oligodendrocytes (OL) [2]. OPCs are activated in response to demyelination [4–6] and altered function and distribution of OPCs have been observed in neurodegenerative [7, 8] and psychiatric diseases [9]. However, difficulties in understanding the ontogenetic complexity, dynamic interactions and functional heterogeneity of OPC populations is a significant barrier to devising OPC based clinical translations.

The origin of OPCs in the spinal cord can be traced back to Olig2-expressing precursor cells that populate the ventral pMN domain and generate vOPCs [10, 11] or Msx3-expressing precursor cells that occupy dorsal ventricular zone domains and generate dOPCs [5, 12–15]. In the spinal cord, the pMN domain generates ~80% of all postnatal OPCs in a sonic hedgehog (Shh) signaling-dependent manner between E12.5 and E14.5 [16–19], while the dorsal Msx3 domain begins to generate the remaining 20% in a Shh-independent manner around E14.5 [12, 13, 20]. OPCs populate the spinal cord in origin specific patterns initially which over time erode by a progressive replacement of early born vOPCs with late born dOPCs [5, 15, 21]. Further, despite the fact that they are derived from different precursor domains, gene profiling- and electrophysiological-analyses suggests that dOPCs and vOPCs converge towards similar transcriptional and electrophysiological states in unchallenged animals [15, 22]. Nevertheless, in the context of injury and aging, functional differences between ventral and dorsal OPCs become apparent. For example upon ventral spinal cord demyelination injury in young adult wt animals, dOPCs are preferentially recruited and display enhanced migration, proliferation, and differentiation compared to vOPCs [5, 6]. These observations leave unexplained the purpose of the sequential production of ventral and dorsal OPC and their possible unique functions remain ill defined.

OPC lineage specific functions might become more readily apparent by the selective ablation of the phylogenetic and ontogenetic older vOPCs since early born OPCs might influence development and distribution of later born OPCs. However, the selective ablation of vOPCs in the spinal cord has so far not been achieved without also inhibiting the earlier production of motor neurons (MN) from the same precursor domain, causing embryonic lethality, and curtailing functional analysis in the postnatal spinal cord [12, 14, 23]. We recently found that the partial and conditional genetic ablation of Shh expression from ventral ventricular zone derivatives in Olig2-Cre expressing embryos (Shh_Olig2^−/−^_) results in a severe reduction in vOPC production without affecting motor neuron (MN) generation during development (Starikov L. and A.H. Kottmann (2020) “Multiple sources of Shh are critical for the generation and scaling of ventral spinal cord oligodendrocyte precursor populations”. Preprint BioRXiv; https://doi.org/10.1101/534750). Here we found that these Shh_Olig2^−/−^_ mice are born alive and mobile and provide evidence for the ectopic expansion of dOPCs throughout their spinal cords. In contrast to OPCs in controls, OPCs in mutants display a distinct morphology and are unable to participate in synaptic remodeling of MNs in response to MN injury. OPCs in the vicinity of degenerating MNs in the SOD1 model of ALS exhibit a morphology reminiscent of OPCs in Shh_Olig2^−/−^_ mice suggesting that compensatory proliferation of dOPCs in response to the degeneration of vOPCs in ALS might result in aberrant morphology and dysfunction in the remaining OPCs.

## Results

### Compensatory expansion of dOPC in embryos with reduced vOPC production

We previously found that lowering of Shh signaling strength in the ventral neural tube achieved by the ablation of a conditional Shh allele (Shh^C/C^; [24]) by Olig2-Cre (abbreviated as Shh_Olig2^−/−^_) from midline derivatives (mainly lateral floorplate and nascent motor neurons, MNs), results in a severe reduction in the numbers of vOPCs that are normally generated from E12.5-E14.5 (Starikov and Kottmann, 2019; BioRxiv). During this time period we observed an almost complete exhaustion of ventricular zone precursor cells within the pMN domain due to a failure of precursor cell maintenance resulting in a 7-10 fold reduction in vOPCs production without reducing motor neuron numbers (Starikov and Kottmann, 2019; BioRxiv). In contrast to other genetic manipulations that curtailed vOPC generation and led to embryonic lethality [12, 14, 23], Shh_Olig2^−/−^_ mice are born alive, morphologically inconspicuous, and mobile, but die around postnatal day P20 (Fig. S1). However, staining of spinal cord cross sections with the OPC marker NG2 (also known as chondroitin sulfate proteoglycan 4) suggested a loss of fine arborization, swelling of soma, and disturbed tiling of OPC in Shh_Olig2^−/−^_ pups (**Fig. 1 A**). Surprisingly, marking OPC somas and quantification of OPC density in Shh_Olig2^−/−^_ mice at P20 in the gray matter of the ventral horns (VH) where vOPCs normally reside [15]), reveals a 29% (+/− 8 %) increased density of OPCs in the vicinity of MNs compared to Shh^C/C^ controls (**Fig. 1 B, C, E**). We did not detect any difference in the numbers of MNs (**Fig. 1 D**) or in the density of NG2^+^ OPCs in the dorsal funiculus (DF), which is mainly populated by dOPCs [15](**Fig. 1A, B, F**).

**Figure 1.**
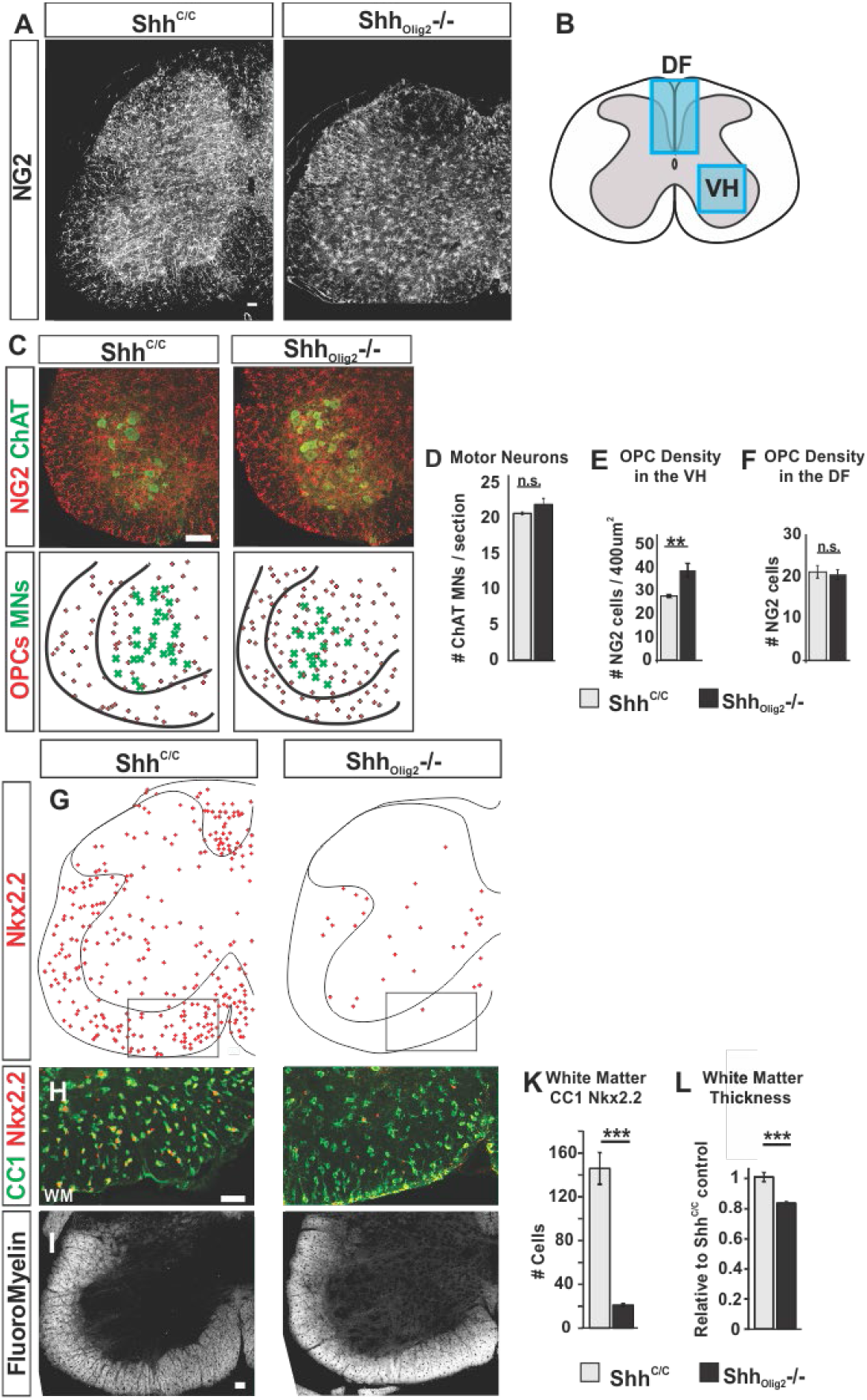
Increased OPC density but depletion of ventrally derived Nkx2.2^+^ oligodendrocytes in Shh_Olig2^−/−^_ P20 pups. (**A**) NG2 staining on lumbar spinal cord hemi sections. Scale bars: 50 μm. (**B**) Schematic depiction of spinal cord areas analyzed, dorsal funiculus (DF), ventral gray matter (GM). (**C**) Staining of MNs and OPCs and derived localization of the soma center of MNs (green X’s) and OPCs (red dots) in ventral horn GM. Scale bars: 50 μm. (**D**) Quantification of NG2+ OPCs density in ventral horn GM. Shh^C/C^ (n = 7 mice), Shh_Olig2^−/−^_ (n = 6 mice). Means ± SEM are shown; Student’s t test. ∗∗p < 0.01. (**E**) Quantification of MNs on L4-L5 lumbar spinal cord hemi sections. Shh^C/C^ (n = 4 mice) and Shh_Olig2^−/−^_ (n = 5 mice). Means ± SEM are shown, Student’s t test. (**F**) Quantification of NG2^+^ OPCs in DF. Shh^C/C^ (n = 3 mice), Shh_Olig2^−/−^_ (n = 6 mice). Means ± SEM are shown, Student’s t test. (**G**) Representative tracing of Nkx2.2^+^ cells on lumbar hemi sections. (**H**) Co-localization of CC1 and Nkx2.2 in ventral white matter OLs. Scale bar: 50 μm. (**I**) Fluoromyelin staining of ventral white matter. Scale bar: 50 μm. (**K**) Quantification of average total white matter Nkx2.2^+^ CC1^+^ OL population. (**L**) Quantification of ventral horn white matter thickness relative to Shh^C/C^ control. (**K,L**) Shh^C/C^ (n = 4-5 mice), Shh_Olig2^−/−^_ (n = 4-5 mice). Means ± SEM are shown, Student’s t test. ∗p < 0.05, ∗∗p < 0.01, ∗∗∗p < 0.001.

We considered two mechanisms that could result in an increased OPC density in the ventral horns despite severely diminished embryonic vOPC production: (1) The few vOPCs that emerge from the pMN domain in mutants disperse and proliferate at a higher rate or for a prolonged period resulting in increased numbers of vOPCs eventually. (2) dOPC production is increased in response to reduced numbers of vOPC and the ventral horns become populated by ectopic dOPC that fail to tile properly.

To distinguish these possibilities, we first quantified the prevalence of oligodendrocytes (OL) that develop from vOPC in the spinal cord of mutant and control animals at P20. vOPC and dOPC lineages can be distinguished in the same animal by genetic tracing in Msx3-Cre; Sox10^GFP/tdTomato^ mice in which dorsally-derived OLs are labeled with tdTomato and ventrally-derived OL with GFP[15]; Fig. S2A). Using this strategy, and as reported before, we find that dOLs preferentially occupy the dorsal and dorsolateral funiculi, while vOLs are found in all remaining regions of the spinal cord (**Fig. S2 B**). Since the dual labeling technique cannot be combined with the conditional ablation of Shh by Olig2-Cre, we utilized Msx3-Cre; Sox10^GFP/tdTomato^ mice to first validate potential molecular markers of ventral oligodendroglial lineage derivatives in the postnatal spinal cord. One such candidate marker for vOL is Nkx2.2. Co-expression of Olig2 and Nkx2.2, a high threshold target gene of Shh signaling in the embryonic spinal cord, was previously found to mark a subpopulation of ventral pMN precursor cells, termed p*-domain, during OPC production [5, 14, 25] [23, 26]. The p* domain is severely reduced in Olig2_Shh^−/−^_ mutants (Starikov and Kottmann, 2019; BioRxiv). Nkx2.2 expression is also found in a subpopulation of white matter (WM) vOLs at E15.5 [23] suggesting that Nkx2.2 could potentially be a marker for the ventral oligodendroglia lineage in the developing postnatal spinal cord as well. Confirming that Nkx2.2 is a marker for ventral derived OLs at P13, we find Nkx2.2 expression in GFP expressing (ventral derived) OLs but not among tdTomato expressing (dorsal derived) OLs at P13 in Msx3-Cre; Sox10^GFP/tdTomato^ mice in (**Fig. S2 C, D**).

Nkx2.2 remains to be expressed in OL marked by the mature OL marker CC1 (anti-adenomatous polyposis coli clone 1) at P20 (**Fig. 1 G, H**). Quantification of the numbers of Nkx2.2^+^ OLs revealed a ~7-fold reduction in Shh_Olig2^−/−^_ mutants compared to controls in the ventral WM at P20 (**Fig. 1 H and K**). Consistent with reduced numbers of mature vOLs, we find decreased WM thickness of ~20% (+/− 4%) in Shh_Olig2^−/−^_ mice, compared to Shh^C/C^ litter controls (**Fig. 1 I, L**). Thus, using Nkx2.2 as a validated marker for ventral derived OL in pups, our analysis suggests that the deficit in vOPCs generation in Shh_Olig2^−/−^_ mutants during early embryogenesis does not recover during later development by expansion of remaining vOPCs. We therefore next investigated whether the deficit in the numbers of vOPCs could become over-compensated by increased dOPC production. At E14.5 the dorsal and ventral phases of oligodendrogenesis overlap: in an anterior to posterior progression, the pMN domain -derived vOPC production comes to an end and dOPC production begins (**Fig. 2 A**; [15]). Consistent, we find a posterior to anterior diminishment of Olig2+ pMN cells in controls as they migrate out of the pMN and spread throughout the spinal cord (**Fig. 2 B,** green side bars; quantified in **Fig. 2 C**, light green bars, 3-way ANOVA spinal cord level: F (2, 204) = 48.38 p < 0.0001, followed by Tukey’s multiple comparisons test), indicating that vOPC production is coming to an end at anterior thoracic levels but is still active at lumbar levels. In contrast, in Shh_Olig2^−/−^_ embryos we observe a 2-, 4-, and 12-fold reduction in the numbers of Olig2^+^ pMN cells at anterior thoracic, posterior thoracic, and lumbar levels, resp. (3 way ANOVA, control vs mutant F (1, 204) = 147.4 p < 0.0001; dorsal vs ventral F (1, 204) = 154.3 p < 0.0001, followed by Tukey’s multiple comparisons test; **Fig. 2 B, C**). There were no differences in the numbers of Olig2^+^ cells delaminating from the dorsal Msx3 domain at anterior thoracic levels (red bars in **Fig. 2 B, C**) and very few to none delaminating Olig2^+^ cells could be detected at posterior thoracic and lumbar levels, resp. in both controls and mutants (**Fig. 2 B,** red side bar, **Fig. 2 C,** arrows). Thus, the quantification of Olig2^+^ cells in the pMN domain and of nascent Olig2^+^ cells emerging from the Msx3 domain suggested that ventral OPC production is severely diminished while dOPC production is initiated and progresses undisturbed in Shh_Olig2^−/−^_ embryos.

**Figure 2.**
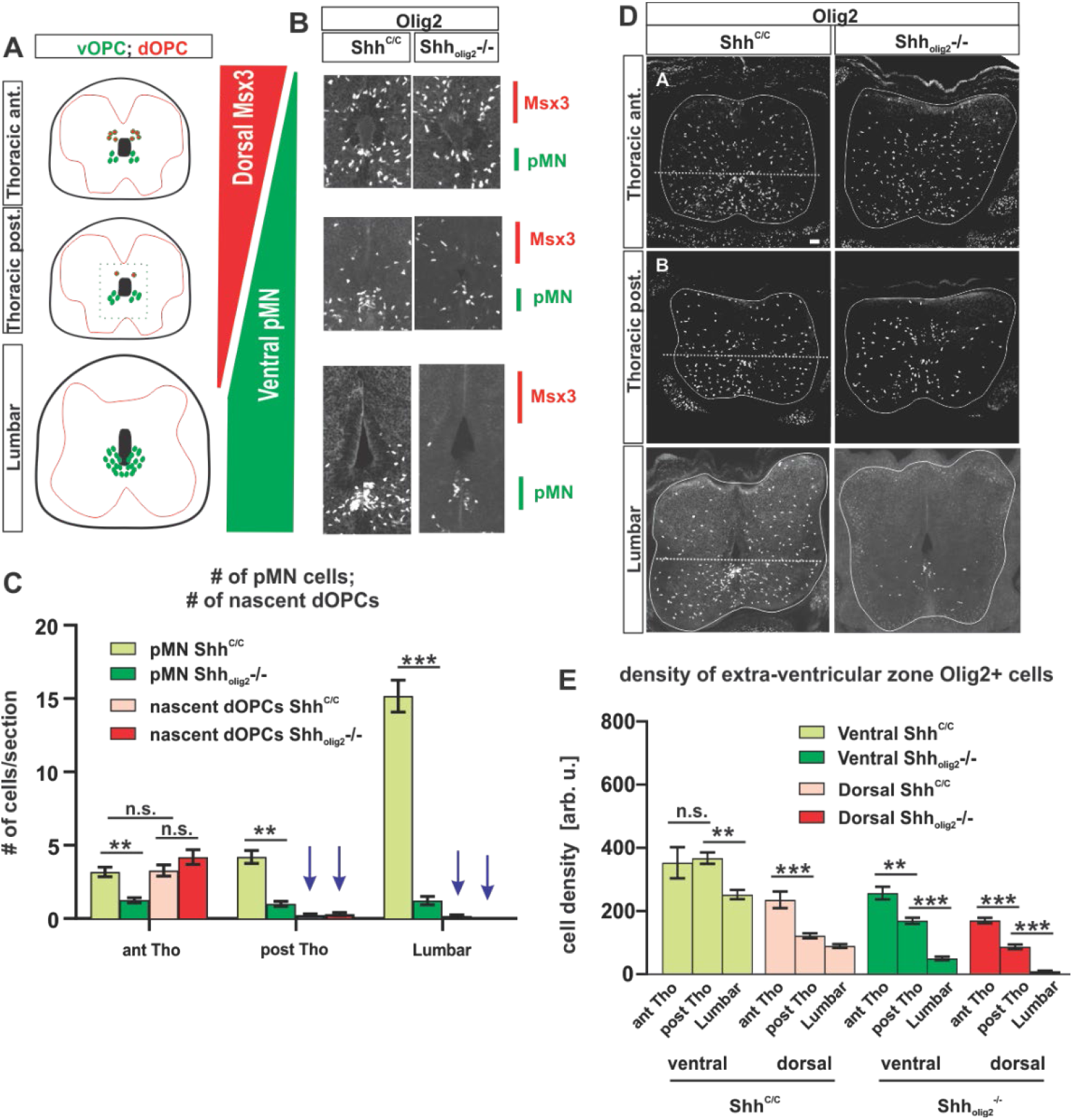
Shh_Olig2^−/−^_ spinal cords become populated by dOPC. (A) schematic depiction of dorsal and ventral OPC precursor domain activity at E14.5. red dots and red bars: dorsal OPC precursor domain; green dots and green bars: ventral OPC precursor domain. (B) Olig2^+^ cells along the ventricular zone at thoracic and lumbar levels of Shh^C/C^ and Shh_Olig2^−/−^_ embryos. Red and green side bars depict approximate location of dorsal and ventral OPC precursor domains within the ventricular zone. (C) Quantification of numbers of Olig2^+^ cells in the ventral pMN domain (green bars) and adjacent to the dorsal ventricular zone (red bars). N=3 per genotype, 3-way ANOVA followed by Tukey’s multiple comparisons test, ∗p < 0.05, ∗∗p < 0.01, ∗∗∗p < 0.001. (D) Immunostaining for Olig2^+^/PDGFRα^+^ OPCs on anterior thoracic, posterior thoracic, and lumbar sections Shh^C/C^ and Shh_Olig2^−/−^_ embryos at E14.5. dotted line above dorsal of pMN domain dissects spinal cord into dorsal and ventral parts. Scale bar: 50 μm. (E) Quantification of OPC density in dorsal (pink/red bars) and in ventral (light green/green bars) spinal cord. N=3 embryos, 3-way ANOVA followed by Tukey’s multiple comparisons test, ∗p < 0.05, ∗∗p < 0.01, ∗∗∗p < 0.001.

We next compared the density of OPCs that have exited the ventricular zone domains (pMN and Msx3) along the spinal cord at E14.5 in mutants and controls to determine if populating of the spinal cord with OPCs in Shh_Olig2^−/−^_ embryos coincides with the activation of dorsal OPC generation. Consistent with the posterior to anterior diminishment of Olig2+ pMN cells in controls at E14.5 (**Fig. 2 C**), we find in the ventral spinal cord of controls a similar density of OPCs at anterior and posterior thoracic levels but a 30 % reduced density at lumbar levels in suggesting that vOPC genesis at thoracic levels has come to completion while vOPC production at lumbar levels is still active (3-way ANOVA Spinal Cord level: F (2, 202) = 87.59 p < 0.0001, followed by Tukey’s multiple comparisons test; **Fig. 2 D, E**; light green bars). In controls, consistent with the presence of delaminating Msx3 domain Olig2+ cells of anterior thoracic segments (**Fig. 2 C**, pink bar), we find in the dorsal spinal cord a 2-fold greater density of OPCs in anterior thoracic segments compared to posterior thoracic segments indicating that half of the OPCs are of ventral origin and the other half are of dorsal origin (3 way ANOVA, Dorsal vs. Ventral; F (1, 202) = 150.6 p < 0.0001, followed by Tukey’s multiple comparisons test; **Fig. 2 D, E**, pink columns). At lumbar levels in control embryos there are fewer OPCs in the dorsal spinal cord compared to posterior thoracic levels consistent with an absence of dorsal and not yet completed ventral OPC production (**Fig. 2 D, E**; pink bars). In contrast to the situation in controls, OPC density in the ventral and dorsal spinal cord in mutants declines progressively from anterior thoracic to posterior thoracic to lumbar levels (3 way ANOVA, Control vs. Mutant: F (1, 202) = 184.6 p < 0.0001, followed by Tukey’s multiple comparisons test; **Fig. 2 D, E**; green and red bars). At all levels the density of OPCs in mutants is significantly reduced compared to controls. Given the severe diminishment of pMN Olig2^+^ cells (**Fig. 2 C**) the anterior to posterior profile of OPC density in mutants conforms well with an anterior to posterior, and ventral to dorsal, progressive complementation of the diminished numbers of vOPCs by increased production and migration of dOPCs in Shh_Olig2^−/−^_ mutant spinal cords. Together with the 7-fold reduction in Nkx2.2^+^ OLs at P20, these observations suggest that reduced production of vOPCs in Shh_Olig2^−/−^_ mice during early embryogenesis does not recover during later phases of development and instead causes the expansion of the dorsal OPC population leading to a spinal cord which is populated mainly by dOPCs in contrast to controls in which the spinal cord is mainly populated by vOPCs.

### The spinal cord in Olig2_Shh^−/−^_ animals is populated by OPCs with aberrant morphology

A higher OPC packing density could result from altered tiling properties [3]. Consistent with this possibility, imaging across the spinal cord suggested that NG2 expressing OPCs of Shh_Olig2^−/−^_ pups display an aberrant arborization with an apparent loss of distal cellular processes (**Fig. 1A**). We therefore quantified dendritic morphology of well-defined individual OPCs in the ventral horns (VH) of Shh_Olig2^−/−^_ mutants and controls in fixed volumes that covered dendrites up to the 9^th^ order in controls using automated dendrite reconstruction of confocal Z-stack images (**Fig. 3A**). Within these volumes we found that dendrites of OPCs in Shh_Olig2^−/−^_ mice arborize 38% (50μm^2^ ROI, Control 6.9 ± 0.9 vs Mutant 9.5 ± 0.9, dendrite order bin 7-9) more frequently compared to controls (**Fig. 3B**). The volume of proximal dendrites (bins 1-3) was increased by 30% compared to controls (Control 10 ± 0.7μm^3^ vs Mutant 13 ± 0.7μm^3^) (**Fig 3C**). Together, these observations suggested that OPCs in mutant animals display fewer distal- and instead have more voluminous- and more highly branched proximal- processes giving them a stubby and bushy appearance.

**Figure 3.**
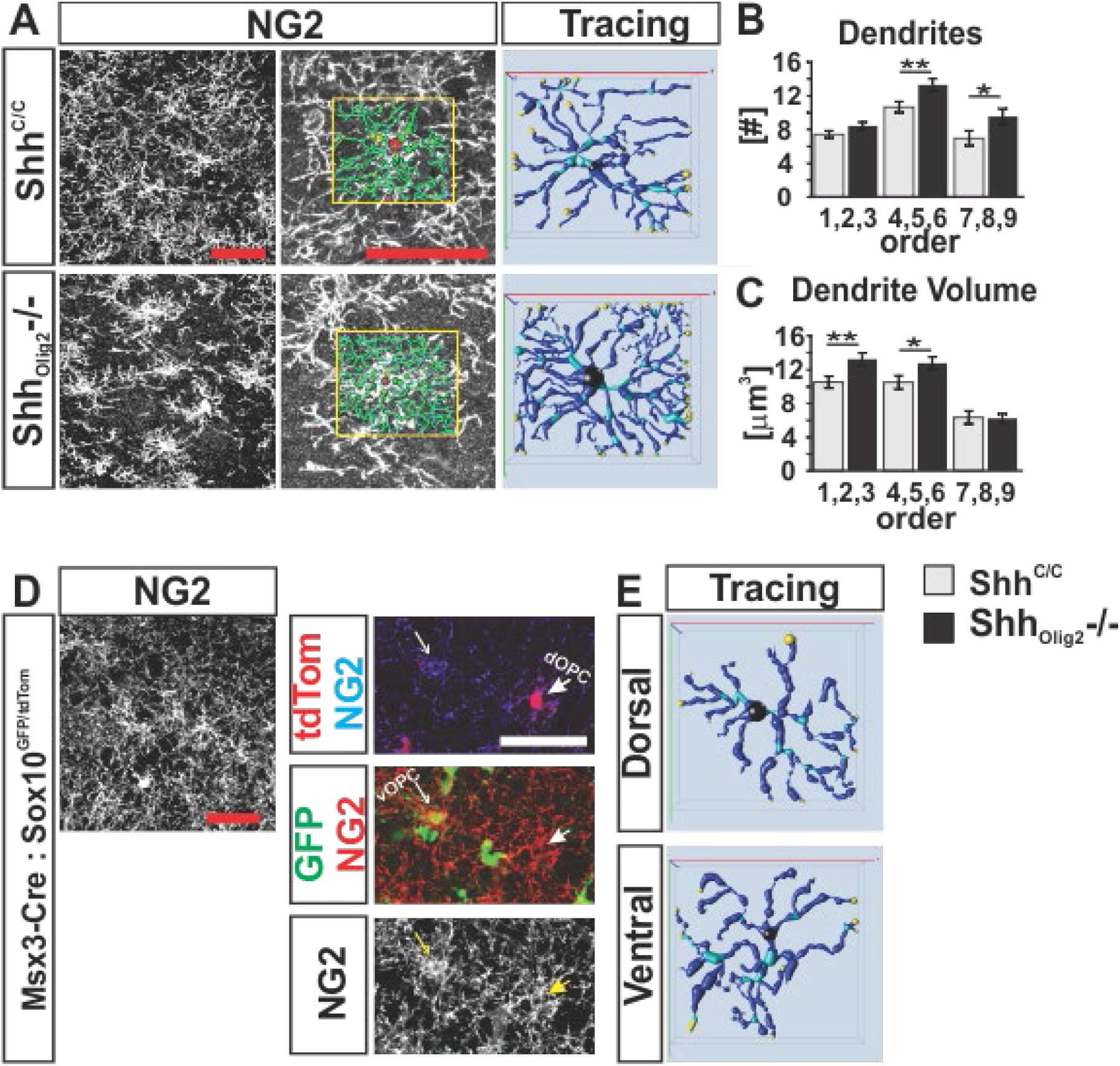
OPCs in mutants display aberrant morphology. (**A**) NG2 stained ventral horn grey matter in Shh^C/C^ and Shh_Olig2^−/−^_ P20 pups. Example of automated tracing and subsequent reconstruction of dendritic arbors of OPCs. Scale bars, 50 μm. (**B**) Quantification of numbers of dendrites per dendritic order. (N=3 mice per genotype, 7-10 cells per animal); Student’s t test. ∗p < 0.05, ∗∗p < 0.01. (**C**) Quantification of dendritic volume per dendritic order. (N=3 mice per genotype, 7-10 cells per animal); Student’s t test. ∗p < 0.05, ∗∗p < 0.01. (**D**) Identification of dOPCs and vOPCs in P13 lumbar spinal cord sections of Msx3-Cre; Sox10^GFP/tdTomato^. (**E**) Reconstruction of dorsal dOPC and ventral vOPC morphology revealed no morphological differences.

We next investigated whether the morphology of dOPC and vOPC differs in wt animals or whether the reduced numbers of vOPCs might induce an aberrant morphology in dOPCs that is distinct from dOPCs in control animals. To distinguish between these possibilities, we analyzed the morphology of cell fate labeled OPCs in P13 lumbar spinal cords of Msx3-Cre; Sox10^GFP/tdTomato^ animals in which dOPCs are labeled by tdTomato and vOPCs by GFP. We found that grey matter dOPCs and vOPCs exhibit a highly similar morphology in wt animals (**Fig. 3D and 3E**), indicating that the unique morphology of dOPCs in Shh_Olig2^−/−^_ is not a distinguishing feature of dOPC in normal animals but rather a consequence of the loss of vOPCs.

### OPCs react towards axotomized motor neurons

OPCs in the ventral spinal cord react to MN denervation from muscle in models of amyotrophic lateral sclerosis (ALS) and are a prominent contributor to gliosis in the spinal cord of ALS patients [7, 27]. Motor nerve injury is used to model aspects of ALS and induces remodeling of selective synaptic terminals on MNs which is thought to prioritize MN recovery over neurotransmission [28–32].

As observed previously [33], 9 days after sciatic nerve axotomy (**Fig. 4A**) we found a pronounced reduction in the numbers of presumptive, excitatory vGlut1 boutons in the immediate vicinity of injured MNs, but not around neighboring, uninjured MNs (**Fig. 4B and 4C**). Conversely, we observed an increased density of cellular processes stained by the OPC marker NG2 around injured MNs compared to uninjured MNs in a pattern that suggested that NG2^+^ cellular processes had taken up space around injured MNs that might be occupied otherwise by vGlut1 labeled boutons on uninjured MNs (**Fig. 4D and 4E**).

**Figure 4.**
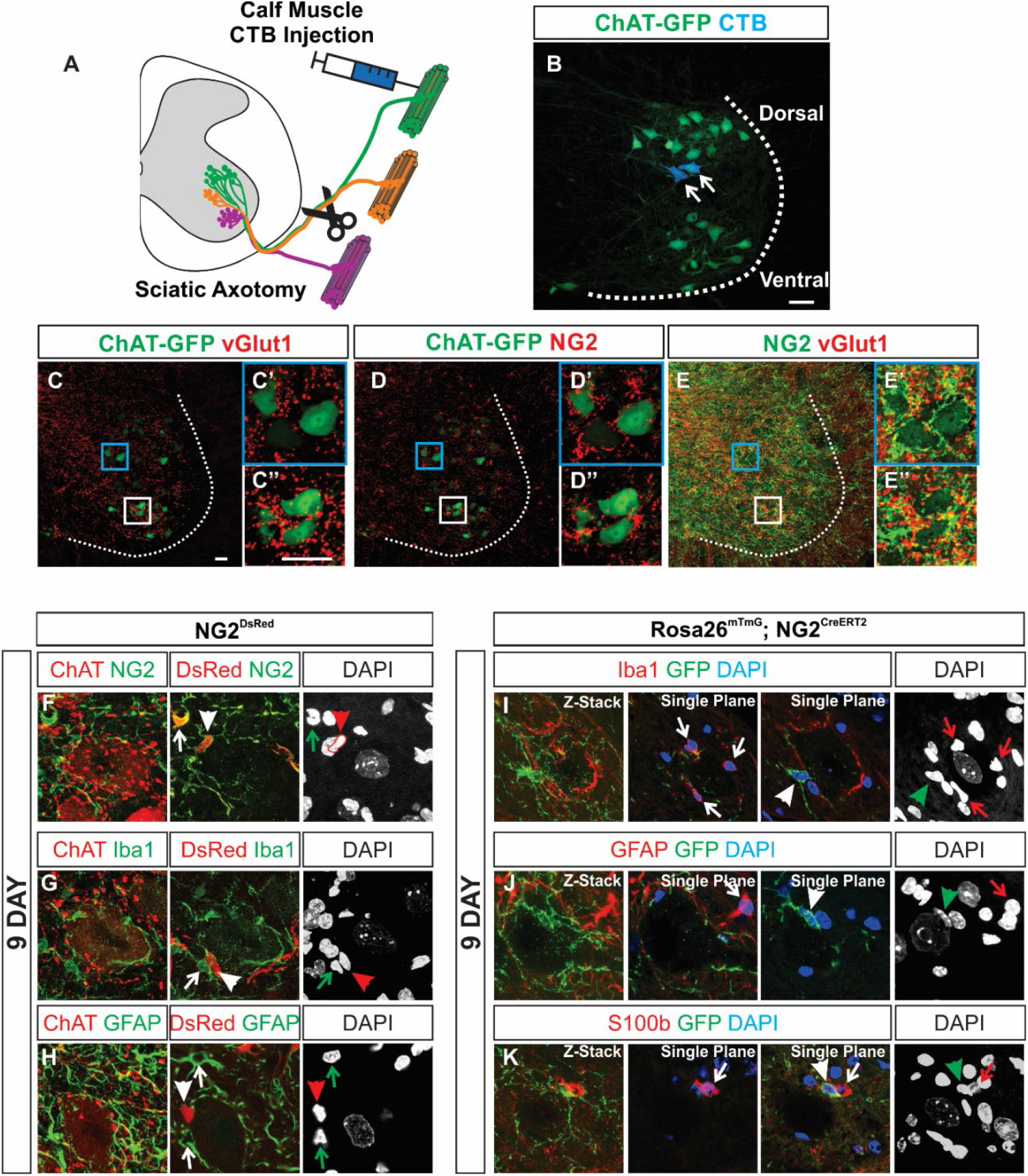
OPCs enwrap axotomized MNs. (**A**) Schematic depiction of CTB labeling and axotomy of MN pools contributing to the sciatic nerve. (**B**) CTB labeled MNs. (**C-E**) Immunostaining with (**C**) vGlut1, (**D**) NG2, (**E**) merge, of ventral horns 9 days post axotomy. (**C’-E’**) Dorsal axotomized MNs are selectively enwrapped by NG2 processes and lose vGlut1 synapses compared to (**C’’-E’’**) ventral uninjured MNs. (**F-H**) Double labeling in NG2^DsRed^ mice 9 days post axotomy with NG2, Iba1, and GFAP. NG2^DsRed^ is expressed in OPCs (arrowhead) and pericytes (arrow) (**F**) NG2+ OPCs reacting and wrapping axotomized ChAT+ MNs, are double labeled with DsRed. (**G**) Iba1 microglia reacting to injured MNs are DsRed negative. (**H**) GFAP astrocytes reacting to injured MNs are DsRed negative. (**I-K**) Double labeling with Rosa26^mTmG^; NG2^CreERT2^ induced mice 9 days post axotomy. GFP labeled cells located on MN somas in close proximity with Iba1, GFAP, and S100b expressing cell somas and processes. GFP OPCs (arrowhead), Iba1 microglia, GFAP and s100b astrocytes (arrows). (**l**) Iba1 microglia reacting to injured MNs are GFP negative. (**J**) GFAP astrocytes reacting to injured MNs are GFP negative. (**K**) s100b astrocytes reacting to injured MNs are GFP negative.

We next investigated whether NG2 positive processes that appeared to enwrap injured MNs could belong to reactive microglia or astrocytes both of which are implicated in protective synaptic pruning [34–36]. We first double labeled spinal cord sections of NG2^DsRed^ expression tracer animals for OPC, microglial, and astrocyte markers. DsRed cell somas were found contacting injured MN somas and stained positive for NG2 (**Fig. 4F**). We did not find double labeling of DsRed together with microglial marker Iba1 or astrocyte marker GFAP (**Fig. 4G and 4H**). We next tested if NG2 expression could be induced acutely in non-OPCs using Tamoxifen induced Cre activity expressed from the NG2 locus. Here we crossed NG2^CreERT2^ with the reporter allele Rosa26^mT/mG^, Tamoxifen was administered intraperitoneally daily for five consecutive days starting on day 4 post axotomy and spinal cords were harvested on day 9 (**Fig. S3A**). Five days of induction resulted in labeling throughout the spinal cord white and gray matter (**Fig. S3B)**. GFP labeled cells had morphology consistent with OPCs or mature oligodendrocytes (**Fig. S3C and S3D**). GFP labeled cells were found wrapping injured MNs (**Fig. 4I-K**). Remarkably, GFP labeled cell somas were often found contacting microglia and astrocyte somas, which were all commonly found in contact with injured MN somas, however, we found no double labeling of GFP with microglial marker Iba1 (**Fig. 4I**) or astrocyte markers GFAP (**Fig. 4J**) and S100b (**Fig. 4K**). These studies indicate that NG2 is neither constitutively expressed nor acutely induced upon injury in microglia or astrocytes. In contrast, we find NG2 being expressed by fate mapped OPCs which enwrap the soma of injured MNs.

We next determined the sequence by which OPCs react towards injured MNs in relationship to other glia subtypes. 3 days post axotomy, we could identify reactive Iba1^+^ microglia and GFAP^+^ astrocytes (**Fig. S4A and S4B**) surrounding injured MNs. At this time however, we did not observe the down-regulation of ChAT, a marker for MN physiological cell stress, loss of vGlut1 synapses, or enwrapment of MNs by NG2 cells (**Fig. S4C and S4D**). However, 9 days post axotomy revealed continued microglial and astrocyte reactivity (**Fig. S4E and S4F**), down-regulation of ChAT in MNs, and the selective enwrapment of injured MNs by NG2 cells concomitantly with the loss of vGlut1 synapses (**Fig. S4G and S4H)**. Thus, the temporal pattern of glia subtype activation indicates that NG2^+^ OPCs respond to MN injury subsequent to microglial and astroglial activation. Together these observations revealed that NG2 expressing OPC react towards axotomized MNs and suggested that OPCs might participate in their synaptic remodeling.

### Injured MNs in Shh_Olig2^−/−^_ are not wrapped by OPCs and are not efficiently stripped of vGlut1 synapses

We next investigated whether OPCs in Shh_Olig2^−/−^_mutants would participate in axotomy induced synaptic remodeling. Since Shh has been shown to be expressed and up-regulated in axotomized MNs [37] Starikov and Kottmann 2019; BioRxiv, and Olig2-Cre activity leads to ablation of Shh from nascent MNs (Starikov and Kottmann, 2019; BioRxiv), we first determined if Shh expressed by MNs would influence the efficacy by which synapses are removed from injured MNs. Shh^C/C^; ChAT-Cre (Shh_ChAT^−/−^_) mutants, like Olig2_Shh^−/−^_ mutants are born with a full complement of motor neurons (**Fig. S5A and S5B**). We next quantified pruning of vGlut1 synapses of injured and non-injured MN somas in Shh^C/C^ controls and Shh_ChAT^−/−^_ mutants. We found that injured MNs are enwrapped by NG2 processes (**Fig. 5A and 5B**) and vGlut1 boutons are eliminated from their surface to the same degree as observed in controls (Shh^C/C^ 34.3% ± 3.6% vs Shh_ChAT^−/−^_ 42.3% ± 1.5%) (**Fig. 5G**), demonstrating that Shh expression by MNs is not critical for these processes.

**Figure 5.**
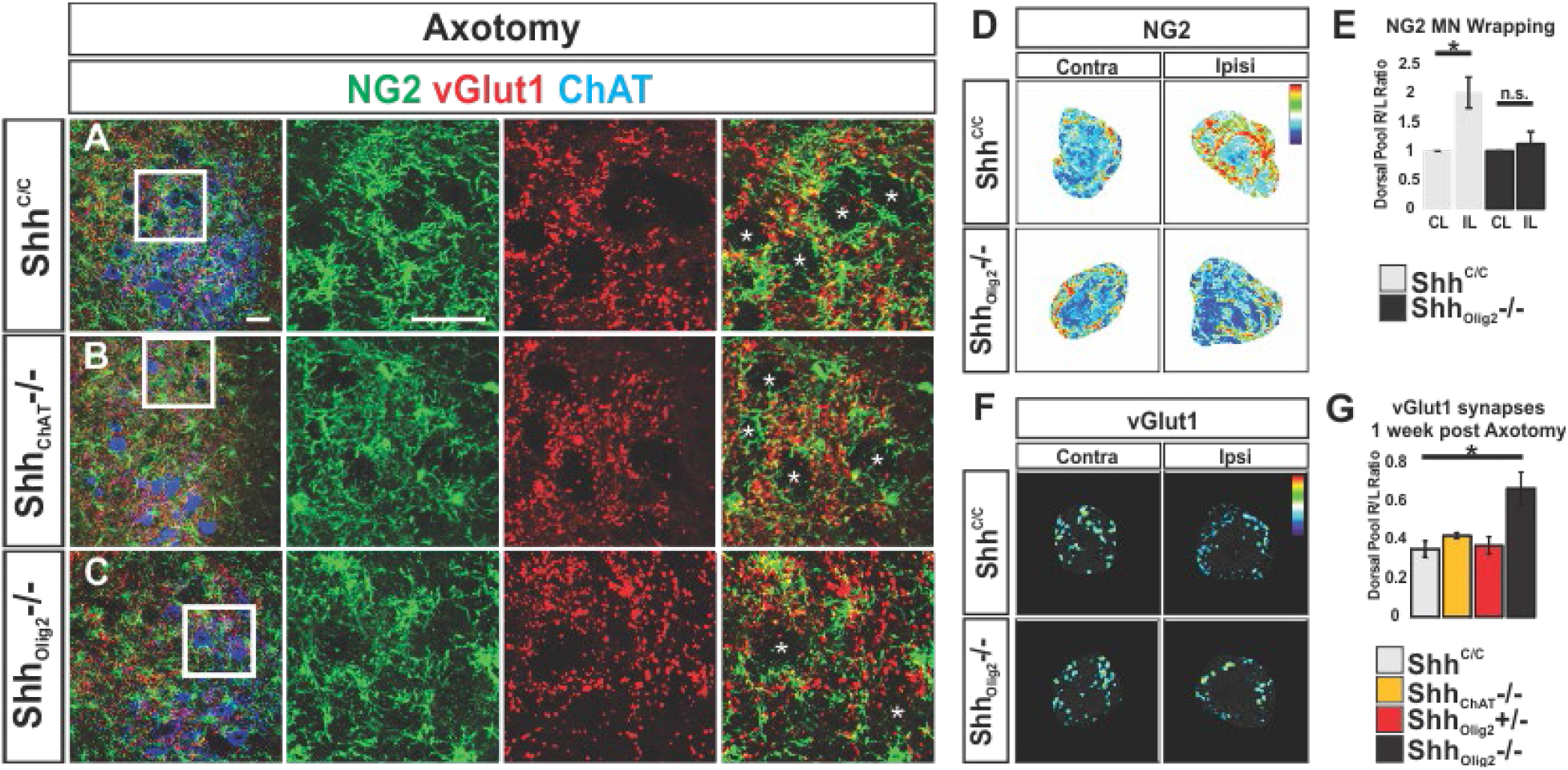
Injured MNs in Olig2_Shh^−/−^_ are not wrapped by dOPCs and are not efficiently stripped of vGlut1 synapses. (**A-C**) Immunostaining of ventral horns with NG2, vGlut1, and ChAT 9 days post axotomy of P20 Shh^C/C^, ChAT_Shh^−/−^_, and Olig2_Shh^−/−^_ mice. Insets of NG2, vGlut1, and merge, asterix indicates MN cell bodies. (**D**) Heatmaps of NG2 processes wrapping and contacting contralateral and ipsilateral MN somas. (**E**) Quantification of contralateral/ipsilateral ratio of NG2 processes contacting MN somas. (**F**) Heatmaps of Ipsilateral and contralateral MN soma stained with vGlut1. (**G**) Contralateral/ipsilateral ratio of surviving vGlut1 synapses on axotomized MN somas. Shh^C/C^ (n = 4 mice), ChAT_Shh^−/−^_ (n = 3 mice), Olig2_Shh^+/−^_ (n = 3 mice), and Olig2_Shh^−/−^_ (n = 3 mice). Means ± SEM are shown. Data were analyzed by Student’s t test. ∗p < 0.05. Scale bars, 50 μm.

In contrast, Olig2_Shh^−/−^_ mutants showed no evidence for enwrapment of injured MNs by NG2 processes (**Fig. 5C, 5D, and 5E**). We analyzed if astrocyte and microglia reactivity around axotomized MNs was affected in Shh_Olig2^−/−^_ mutants and found comparable activation of Iba1 and GFAP cells around axotomized MNs in Shh_Olig2^−/−^_ animals and controls (**Fig. S5C–S5F**). However, consistent with the absence of NG2 processes around injured MNs, we found a preservation of vGlut1 synapses on axotomized MNs when compared to uninjured MNs in Shh_Olig2^−/−^_ mutants (Shh^C/C^ 34.3% ± 3.6% vs Shh_Olig2^−/−^_ 65.5% ± 9.1%) (**Fig. 5F and 5G**). These results indicate that OPCs in Shh_Olig2^−/−^_ mutants cannot participate in efficient pruning of vGlut1 synapses upon motor neuron nerve injury and thus begin to suggest that vOPC production is needed to populate the ventral spinal cord with OPCs that are capable to communicate with injured MNs.

### OPCs in the vicinity of MNs in end-stage SOD1 mutants display altered morphology

Oligodendrocyte dysfunction and death has been observed in ALS [7, 8]. In the SOD1 model of ALS, OPCs dramatically increase their proliferation and differentiation to compensate for the death of dysfunctional mature oligodendrocytes. Since in Shh_Olig2^−/−^_ mutants, dOPCs are forced to over-proliferate to compensate for the absence of vOPC we examined OPC morphology in end-stage (P135) SOD1^G93A^ animals that express the NG2^DsRed^ gene expression allele (**Fig. 6**). In the SOD1 spinal cord we observed OPCs with aberrant morphology which were predominantly localized in the ventral horns (**Fig. 6C-E**) consistent with previous findings that increased OPC proliferation in SOD1 mutants occurs preferentially in the ventral horns near MNs [7]. These NG2^DsRed^ OPCs display swollen soma and swollen primary dendrites compared to aged matched controls (**Fig. 6I and 6J**) giving them a morphology that is reminiscent of OPC morphology in Shh_Olig2^−/−^_ mutants. Our observations suggest that exhaustion of ventral OPCs in ALS might result in proliferation induced morphological changes of recruited dOPCs that consequently might fail to functionally compensate for diminishing numbers of vOPCs.

**Figure 6.**
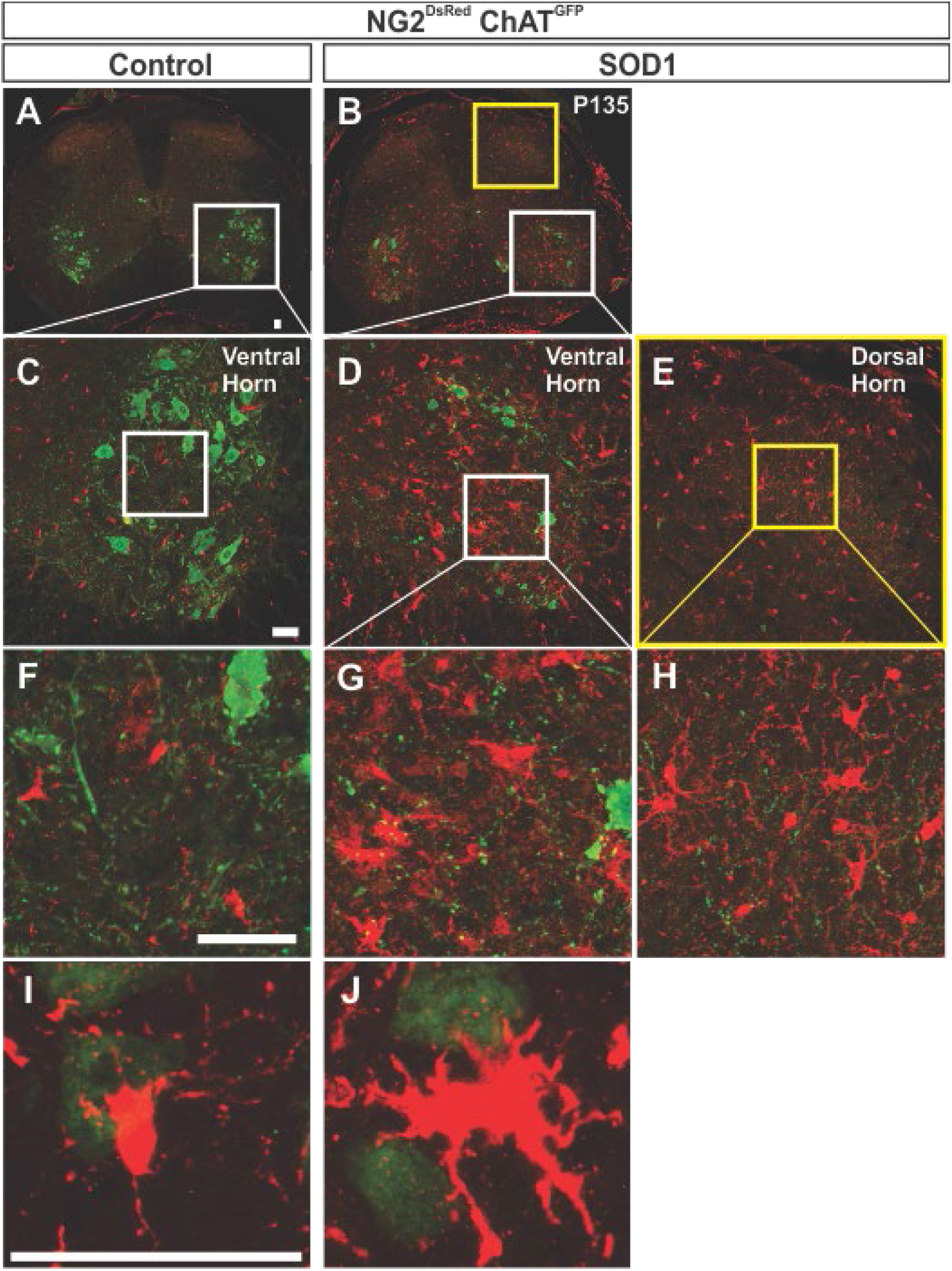
OPCs in end-stage SOD1 mutants display altered morphology. (**A-J**) P135 NG2^DsRed^ ;ChAT^GFP^ control and SOD1 lumbar spinal cord sections. OPCs in end-stage SOD1 animals show higher ramified proximal processes, reminiscent of altered dOPCs morphology in Olig2_Shh^−/−^_ mutants.

## Discussion

In this study we took advantage of the finding that a reduction in Shh signaling strength during early spinal cord development allows the production of a full complement of MNs but results in an almost complete absence of vOPCs (Starikov and Kottmann, Biorxiv 2019). Here we observed that the severely diminished numbers of vOPC in mutant embryos become progressively replenished by increased dOPC production from E14.5 onwards resulting in a greater OPC density compared to controls in the postnatal ventral spinal cord. These OPCs, however, (1) exhibit smaller arbors and tile the parenchyma unevenly; (2) fail to participate in stripping off synapses from injured motor neurons and (3) resemble the morphology of OPCs present in the ventral horns of end stage SOD1 mice. Our data reveal that the earlier production of ventral OPCs prevent later born dorsal OPCs from (1) tiling the spinal cord completely and (2) suggest that vOPCs possess a unique function in responding to MN injury.

### Absence of vOPCs is compensated by expansion of dOPCs of aberrant morphology

Despite the severe reduction in vOPC production, we find that the postnatal ventral spinal cord becomes populated by NG2^+^ OPCs in Shh_Olig2^−/−^_ mutants. Our data conforms well with previous studies demonstrating that dOPCs can be generated in the absence of vOPCs and spread throughout the embryonic spinal cord [12, 14, 23]. The current study extends these earlier observations in two significant ways: The previous approaches for inhibiting vOPC production led to (A) gross disturbances of ventral spinal cord development including an absence of MNs and (B) early embryonic lethality. These experimental shortcomings did not allow to assess the spread and function of dOPCs in the postnatal ventral spinal cord.

We find that the majority of OPCs in the ventral parenchyma of mutants originate from the dorsal Msx3 precursor domain which begins to produce OPCs in a Shh-independent manner at the end of vOPC production in an anterior to posterior wave starting at E14.5 [12–14]. Overall, the result of this compensation is a reversal of the relative distribution of ventral and dorsal OPCs: while the spinal cord in controls is populated by 80 % vOPCs and 20 % dOPC [6], in Shh_Olig2^−/−^_ mutants the majority of OPCs is of dorsal origin. Surprisingly, OPC density in the ventral horns of mutants becomes increased suggesting a disturbance in the regulation of OPC proliferation and/or tiling. OPCs in mutants obtain an aberrant morphology that is distinct from the morphology of vOPC or dOPC in wt animals and is characterized by a loss of distal dendritic branches and, conversely, a thickening of proximal dendrites and a swelling of soma. A parsimonious explanation for the increased density of OPCs in mutants is that a reduced size of their dendritic arbors allows a greater packing density. Thus, our observations reveal that the earlier emergence of vOPCs is critical for preventing excessive proliferation, aberrant morphology and subsequent infiltration into the ventral spinal cord of dOPC.

### Reduced production of vOPCs puts in motion an adaptive process leading to dysfunctional dOPCs that do not react to MN injury

Hyperexcitability of MNs in ALS might contribute to their progressive demise [38]. In the SOD1 mouse model of ALS, ablation of mutant SOD1 selectively from OPCs prolonged the lifespan of animals compared to control SOD1 mice, implicating dysfunctional OPCs in the disease mechanism [7]. Given the age-associated decline in dOPC reactivity to injury [6] and the involvement of OPCs in models of ALS, we investigated if (1) OPCs participate in synaptic remodeling of injured MNs and (2) whether the altered dOPC phenotype we observe in Shh_Olig2^−/−^_ would hinder this process.

Our observations implicate OPCs in synaptic rearrangement in response to peripheral motor nerve injury. We find that the presence of OPC dendrites in apparent close contact with injured MNs is critical for the removal of vGlut1 boutons since we neither observed “wrapping” nor removal of vGlut1 boutons from injured MNs in Shh_Olig2^−/−^_ mice despite the presence of OPCs at greater density in the vicinity of MNs. While OPCs have thus far not been implicated in synaptic pruning as such, OPCs have been found recently to regulate the homeostasis of different microglia cell states [39]. Taken together with previous results demonstrating the involvement of microglia and astrocytes in synaptic pruning our data suggests the sequential involvement of all three glia subtypes in a model in which OPC interaction with MNs is critical for efficient elimination of glutamatergic (vGlut1) synapses following nerve injury.

While our experiments are not able to identify which lineage of OPCs becomes reactive in response to MN axotomy in wt animals, they do provide evidence that dOPCs cannot functionally compensate for the loss of vOPCs in regard of synapse pruning. This finding sheds additional light on how OPC dysfunction might participate in the etiology of ALS: OPCs in end-stage mutant SOD1 animals have been shown to dramatically increase their proliferation rate specifically around dying motor neurons in order to replace degenerating OLs within the ventral horns of the spinal cord [7, 8]. Once the pool of resident vOPCs is exhausted it is thought that proliferative dOPC migrate into the ventral horns. Hence, a commonality between OPCs in Shh_Olig2^−/−^_ and OPCs in SOD1 mutants is increased proliferation. Consistent with this scenario we identified OPCs in the vicinity of degenerating MNs that exhibit a morphology reminiscent of dOPCs in Shh_Olig2^−/−^_ mutants. Our data predicts that these OPCs are unable to contribute to synaptic remodeling of distressed MNs.

More than 20 years ago Tom Jessell remarked that it was “inconceivable” that OL lineages of distinct ontogenetic origins could have completely redundant functions. He further suggested that determining lineage specific functions would require the selective genetic ablation of each OL lineage without disturbing the patterning of the neural tube otherwise. The senior author on this paper, as post doc in Tom’s lab, was unable to design and produce the genetic tools needed at the time. While the current paper begins to confirm Tom’s predictions, the tools presented here are far from perfect and the findings do not amount to what Tom would have classified as a “story” but possibly the beginning of one.

## Acknowledgements

The authors thank Susan Morton (Jessell Lab), Sam Pfaff and Benjamin Novitch for providing antibody reagents, and Richa Tripathi and William Richardson for providing frozen tissue from Msx3-Cre; Sox10^GFP/tdTomato^ pups. The authors thank Dustin Zuelke for help with statistical analysis. The authors thank Dustin Zuelke and Miruna Ghinia-Tegla for many valuable discussions and critical reading of earlier versions of the manuscript. Funding was provided by Research Foundation of the City College of New York to AHK. LS was funded in part by the graduate program in molecular, cellular and developmental biology (MCD) of the graduate center of the City University of New York.

## Materials and Methods

### Transgenic Mice

The following mouse strains were used and genotyped as described previously: Shh-nLZ^C/+^ (Gonzalez-Reyes et al., 2012), Chat-Cre (Rossi et al., 2011), Olig2-Cre (Dessaud et al., 2007), Chat-EGFP/Rpl10a (Doyle et al., 2008), NG2^DsRedBac^ (Zhu et al., 2008), NG2-Cre^ERT2^ (Zhu et al., 2011), Rosa26^mTmG^ (Muzumdar Mandar Deepak et al., 2007), SOD1^G93A^ (Gurney et al., 1994). Mice were maintained on a C57BL/6 background. Mice were kept on a 12 hr dark/light cycle and the day of birth designated P1. All animal experiments were approved by the Institutional Animal Use Care Committee at CUNY.

### Retrograde Motor Neuron Labeling

Animals were sacrificed 1 week after (1ug/ul) Cholera Toxin Subunit B (Recombinant), Alexa Fluor 594 (Thermofisher) was injected in four 2ul injections into the right calf muscle.

### Surgery

For axotomy, the right sciatic nerve was exposed at midthigh level under isoflurane anesthesia and a 2-mm segment of sciatic nerve was removed to avoid regeneration. The wound was closed with silk sutures. Mice were perfused 3 or 9 days after surgery. All surgeries were performed under CCNY IUCUC guidelines.

### Tissue Processing

All mice were sacrificed using an overdose of anesthetic, subjected to transcardial perfusion with 4% (w/v) paraformaldehyde (PFA) in 0.1 M PBS pH 7.4. Spinal cords and embryos were dissected, postfixed in 4% PFA for 1 hr at 4°C, cryoprotected with 30% (w/v) sucrose in 0.1M PBS for 24–48 hr, embedded and frozen in OCT medium, and stored at −80°C. Tissues were sectioned at 20 μm and collected onto glass slides.

### Immunocytochemistry and Microscopy

20um thick spinal cord cryosections were air dried for 30 min. Then sections were washed with PBS for 10 mins and with 0.3% [v/v] Triton X-100 in PBS for 20 min. Sections were then pre-treated with blocking solution (10% [v/v] horse serum and 0.3% [v/v] Triton X-100 in PBS) for 90 mins and incubated with primary antibodies overnight at 4C. The next day, following 3 PBS washes the sections were incubated with secondary antibodies for 2 hr at 20C–25C. A table for all antibodies and reagents used is provided. For cell counts, at least 3 sections per animal from 3 mice were examined, unless otherwise noted. Images were acquired using a Zeiss LSM880 confocal microscope.

Statistical analysis was performed using Prism 7 (Graphpad Software Inc.), two-tailed, paired or unpaired Student’s t test, and two and three way ANOVA followed by Tukey’s multiple comparisons test were used. The data are presented graphically as: ∗(p < 0.05), ∗∗(p < 0.01), and ∗∗∗(p < 0.001).

### Morphological Analysis of OPCs

Images of NG2 stained 20μm thick spinal cord sections were captured with the LSM880 Zeiss confocal microscope using the using oil immersion 40x objective. Z stacks were taken with 0.5μm step intervals, with an average of 35-40 images per stack. 50μm^2^ regions of interest were designated with an OPC soma in the center. Dendrites were reconstructed in 3D using automatic tracing in NeuronStudio (Wearne et al., 2005). Reconstructed OPCs were analyzed with NeuroExplorer software for dendritic order analysis. Data for dendrites were binned in groups of 3. For cell morphology analysis, 7-10 cells were reconstructed per animal from three mutants and controls.

### Analysis of NG2 processes and vGlut1 Terminals on MN Somas

20μm spinal cord sections from P20 lumbar L4-L6 segments with Chat-EGFP MNs were stained with vGlut1 and NG2 antibodies. Dorsal motor pool MNs were outlined in ImageJ in the ChAT-EGFP channel and the outline was expanded by 15 pixels on both contralateral and ipsilateral sides of axotomy. MN outlines were used as regions of interests (ROIs) in vGlut1 and NG2 stained channels. Heatmaps were generated to threshold each image using the Heatmap Histogram plugin for ImageJ. Ratios were generated by comparing contralateral and ipsilateral MNs for each section to control for staining variability between sections.

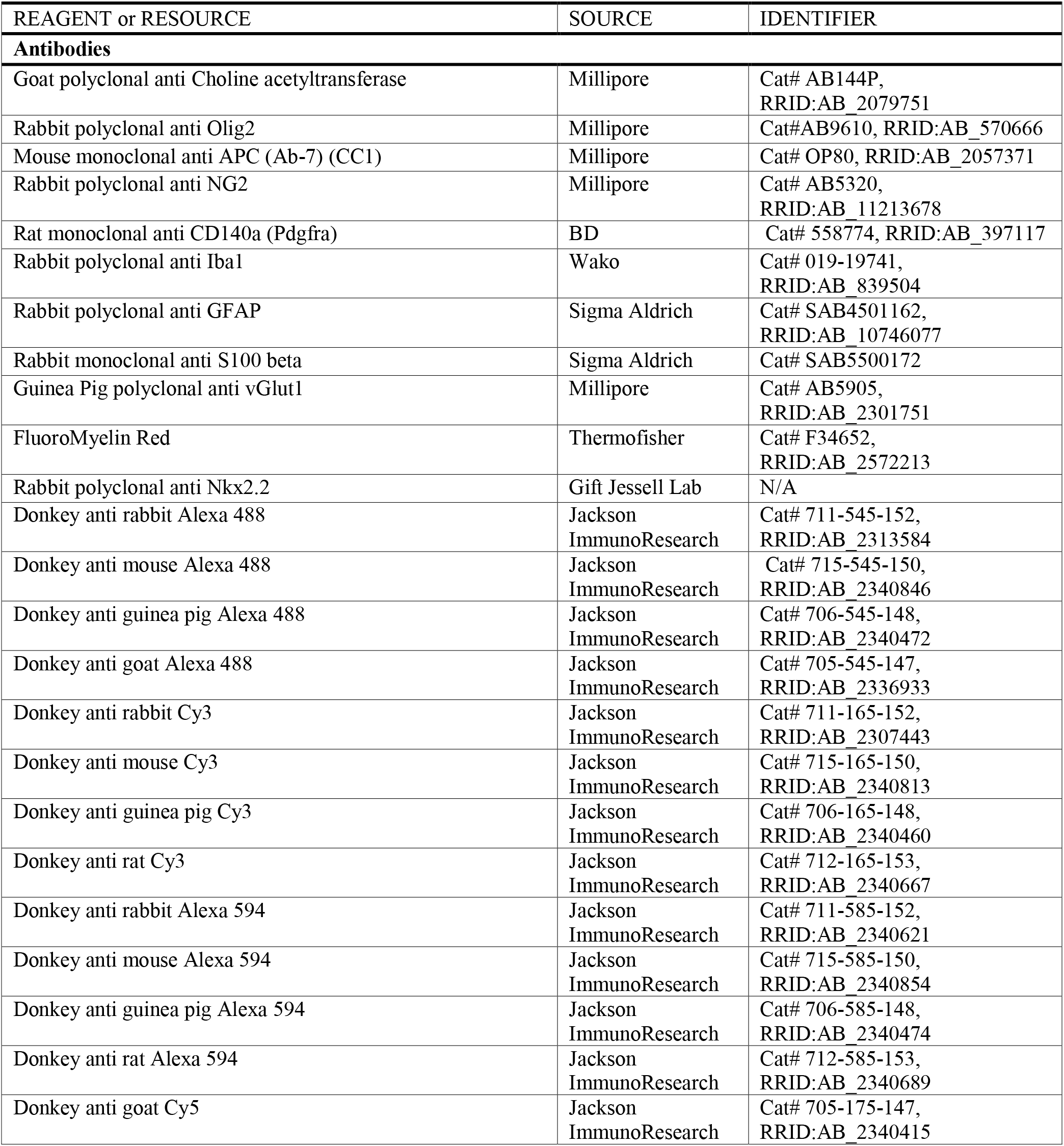
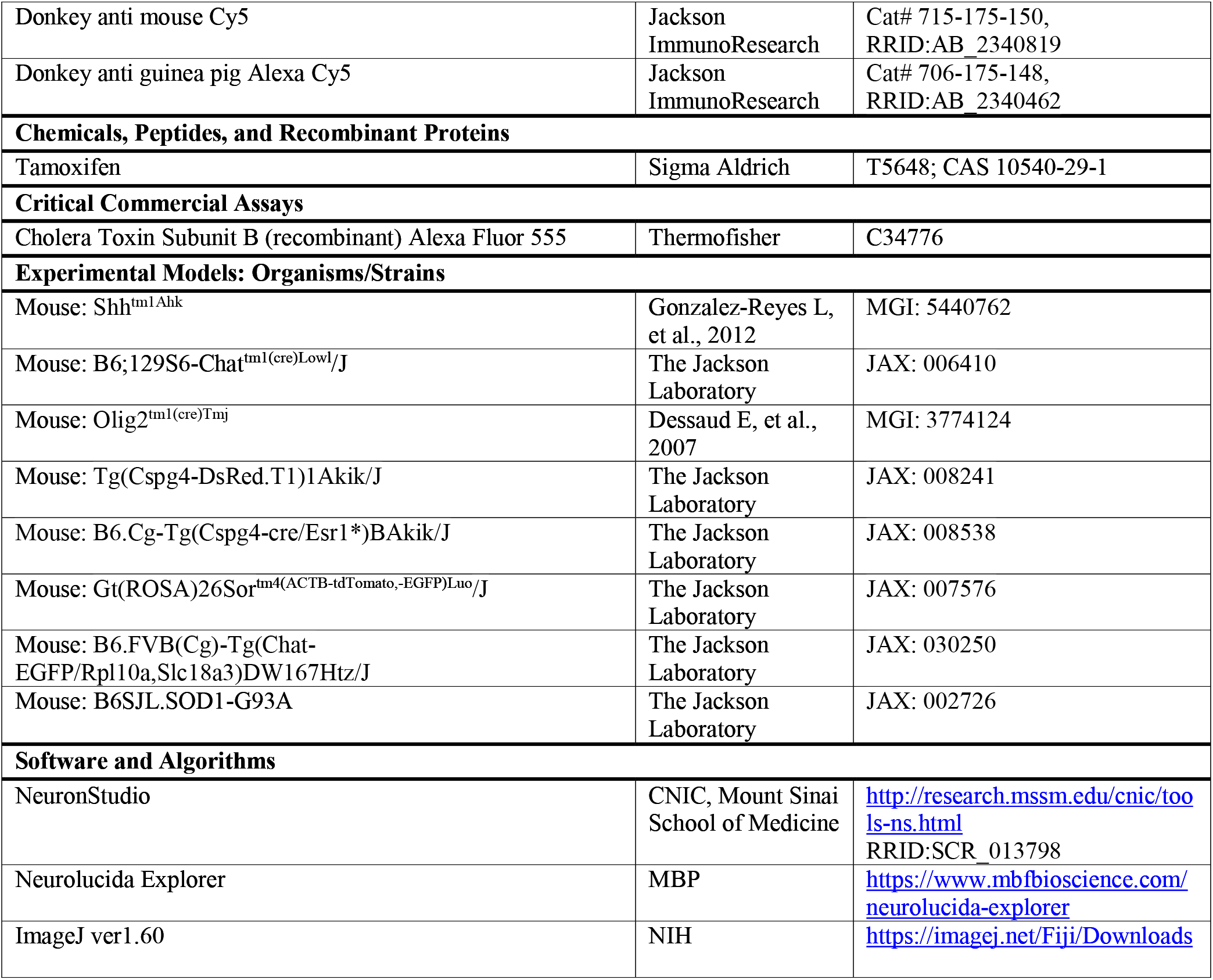

---

**Sup Figure 1.**
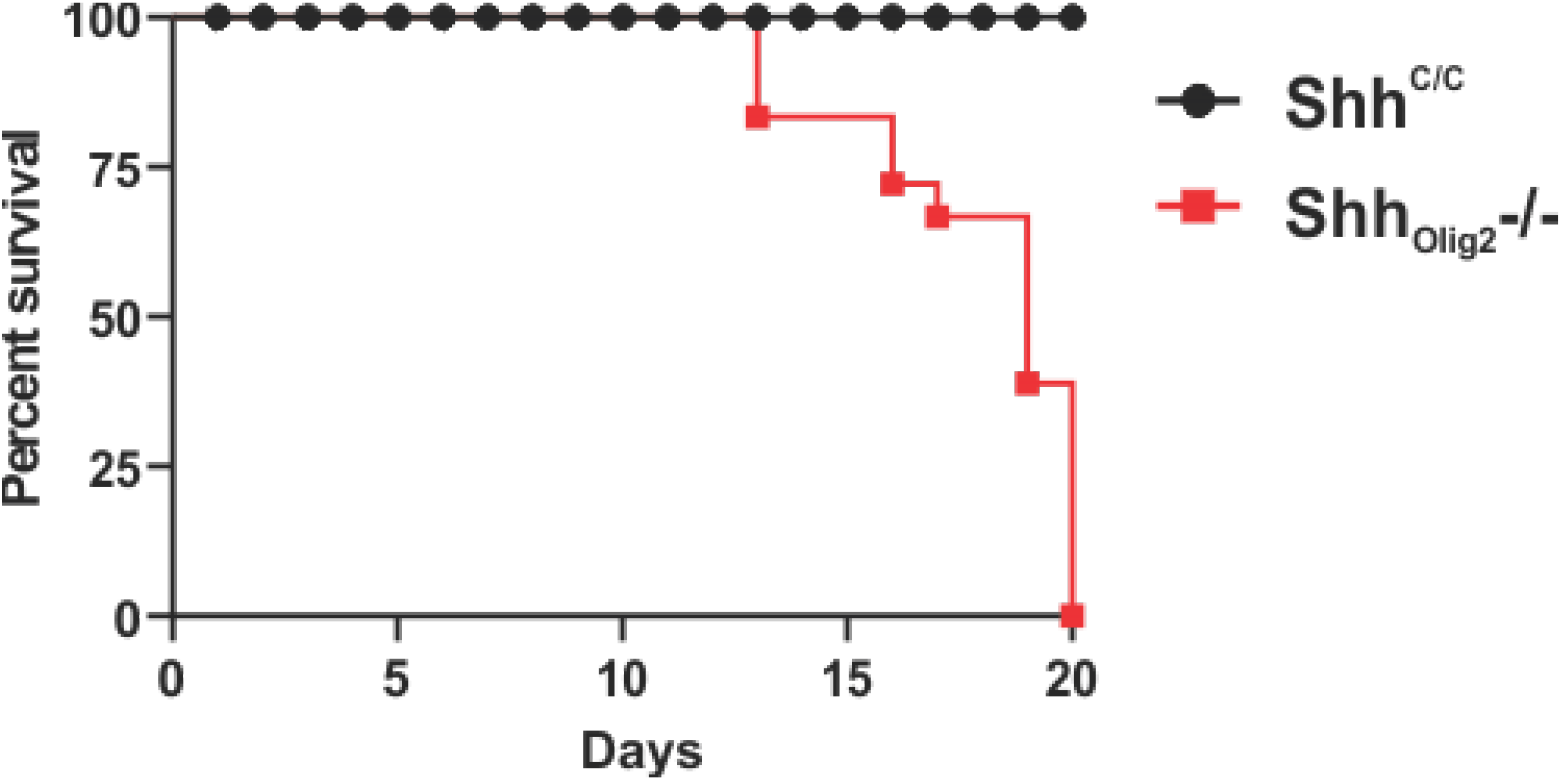
Lifespan of Olig2_Shh^−/−^_ mutants.

**Sup Figure 2.**
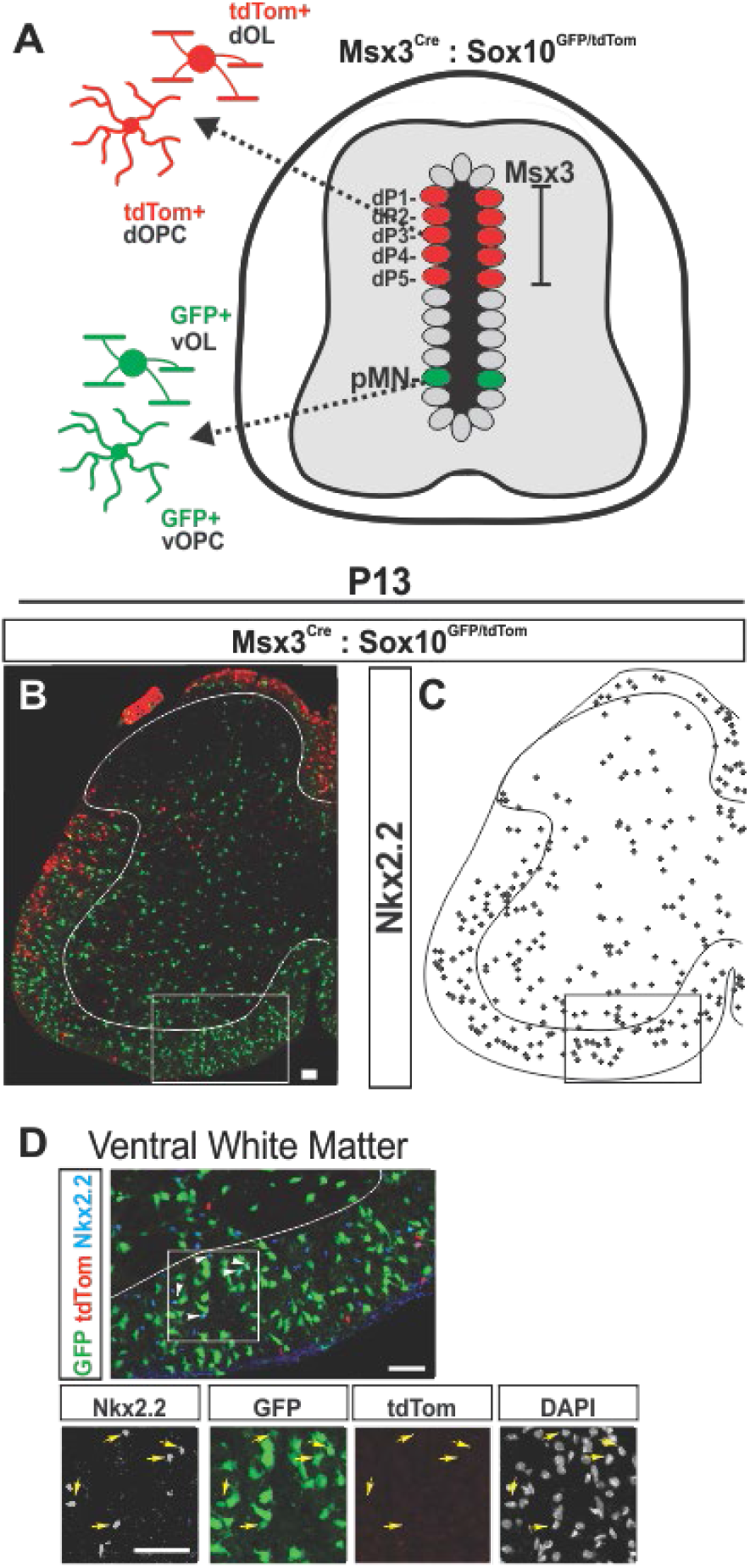
Validation of Nkx2.2 as a postnatal marker of ventral OPCs. (**A**) Schematic of dOPC and dOL production from Msx3 domain and vOPC and vOL production from pMN domain at E14.5. (**B**) Labeling of ventral- and dorsal- oligodendrocyte lineage with Msx3-Cre; Sox10^GFP/tdTomato^. (**C**) Representative tracing of Nkx2.2 expressing cells on P13 Msx3-Cre; Sox10^GFP/tdTomato^ lumbar hemi section. (**D**) Immunolabeling for Nkx2.2 in Msx3-Cre; Sox10^GFP/tdTomato^ ventral white matter. Arrows indicate co-expression of GFP and Nkx2.2 but not tdTomato and Nkx2.2.

**Sup Figure 3.**
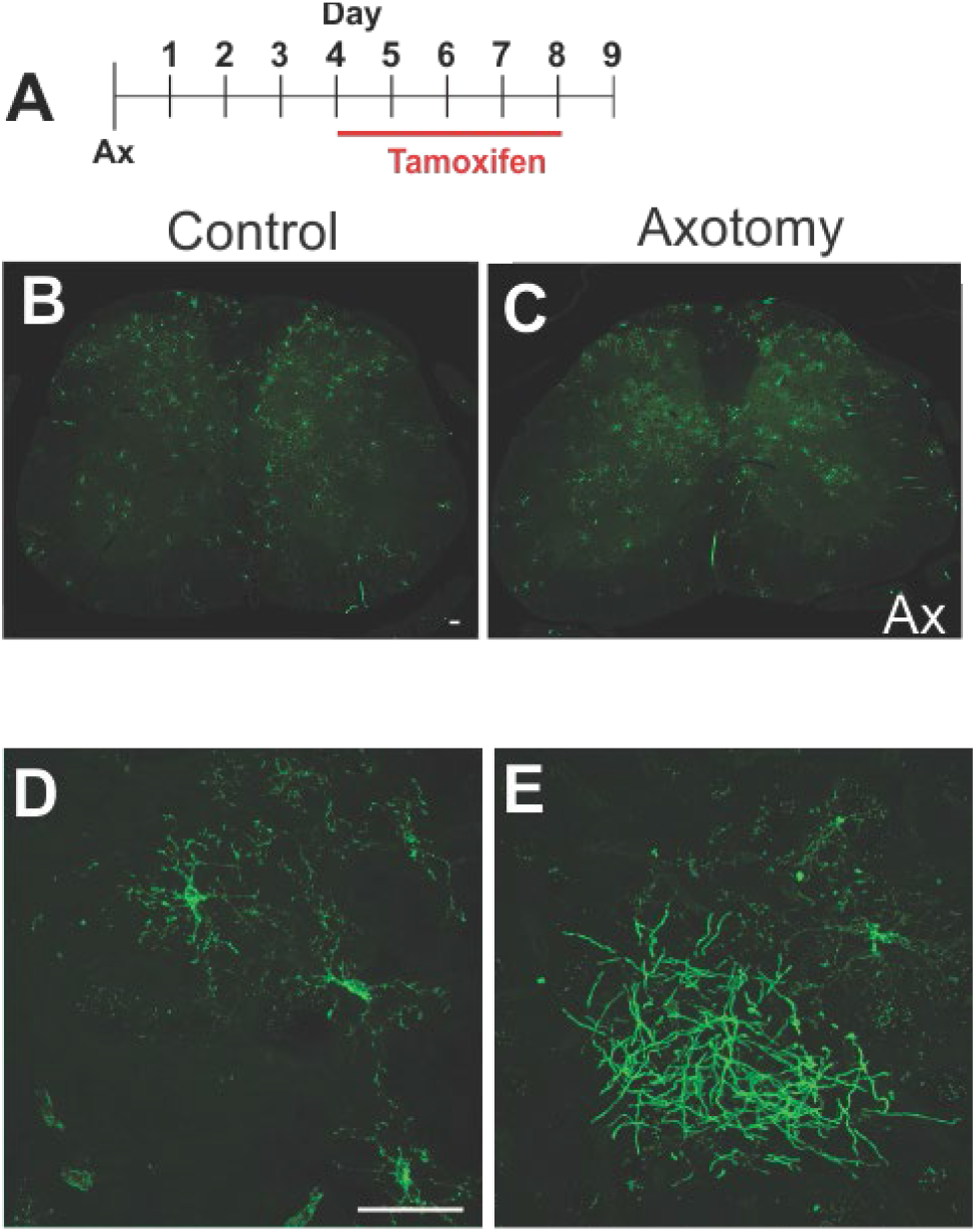
NG2-Cre^ERT2^ selectively labels OPCs and newly differentiated oligodendrocytes. (**A**) Schematic of tamoxifen administration of Rosa26^mT/mG^; NG2-Cre^ERT2^ mice after unilateral axotomy. (**B, C**) mGFP expression marks OPCs and OLs that have differentiated from labeled OPCs throughout spinal cord sections in controls (B) and mice with unilateral axotomy (C). (**C**) mGFP labeled cells displayed OPC morphology (**D**) or mature oligodendrocyte morphology.

**Sup Figure 4.**
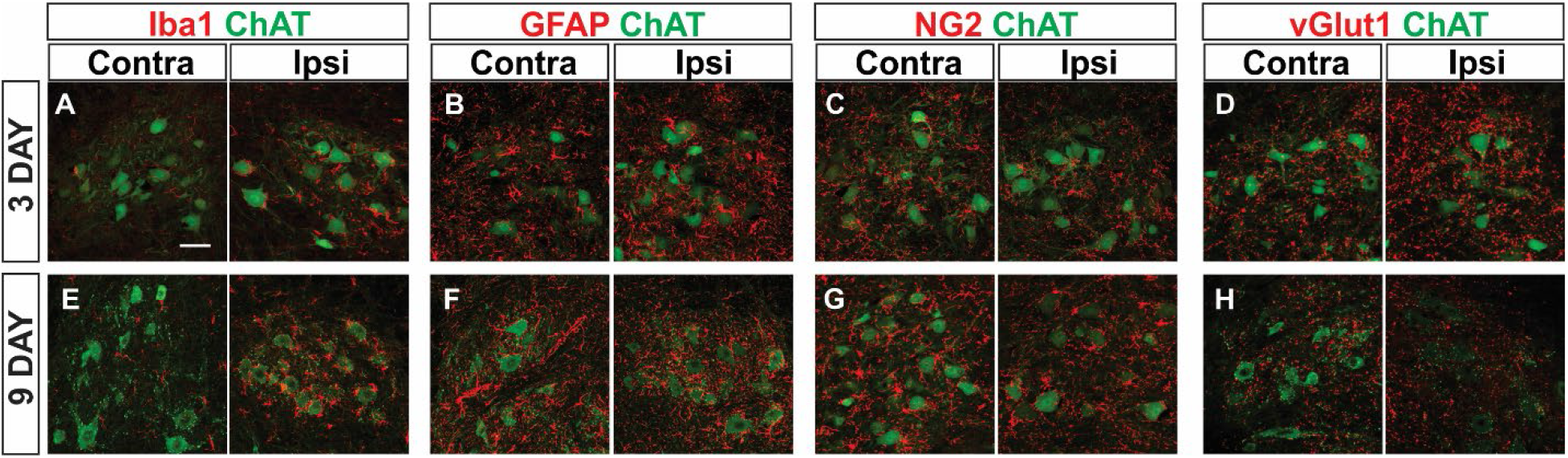
Time course of microglia, astrocyte, and OPC reactivity, and vGlut1 synapse loss in response to axotomy. (**A-D**) Reactive changes in the vicinity of MNs 3 days post sciatic axotomy in control Shh^C/C^. (**A**) Iba1+ microglia and (**B**) GFAP+ astrocytes are reactive and change morphology around injured ipsilateral MNs. (**C**) NG2+ OPCs are not found to be reactive or change morphology around injured MNs. (**D**) vGlut1 terminals are maintained on injured MN somas 3 days post axotomy. (**E-H**) Reactive changes in the vicinity of MNs 9 days post sciatic axotomy. (**E**) Iba1+ Microglia and (**F**) GFAP+ astrocytes remain reactive, in addition to the (**G**) selective enwrapment of injured MNs by NG2 processes as well as (**H**) a loss of vGlut1 terminals on MN somas and down-regulation of ChAT in injured MNs.

**Sup Figure 5.**
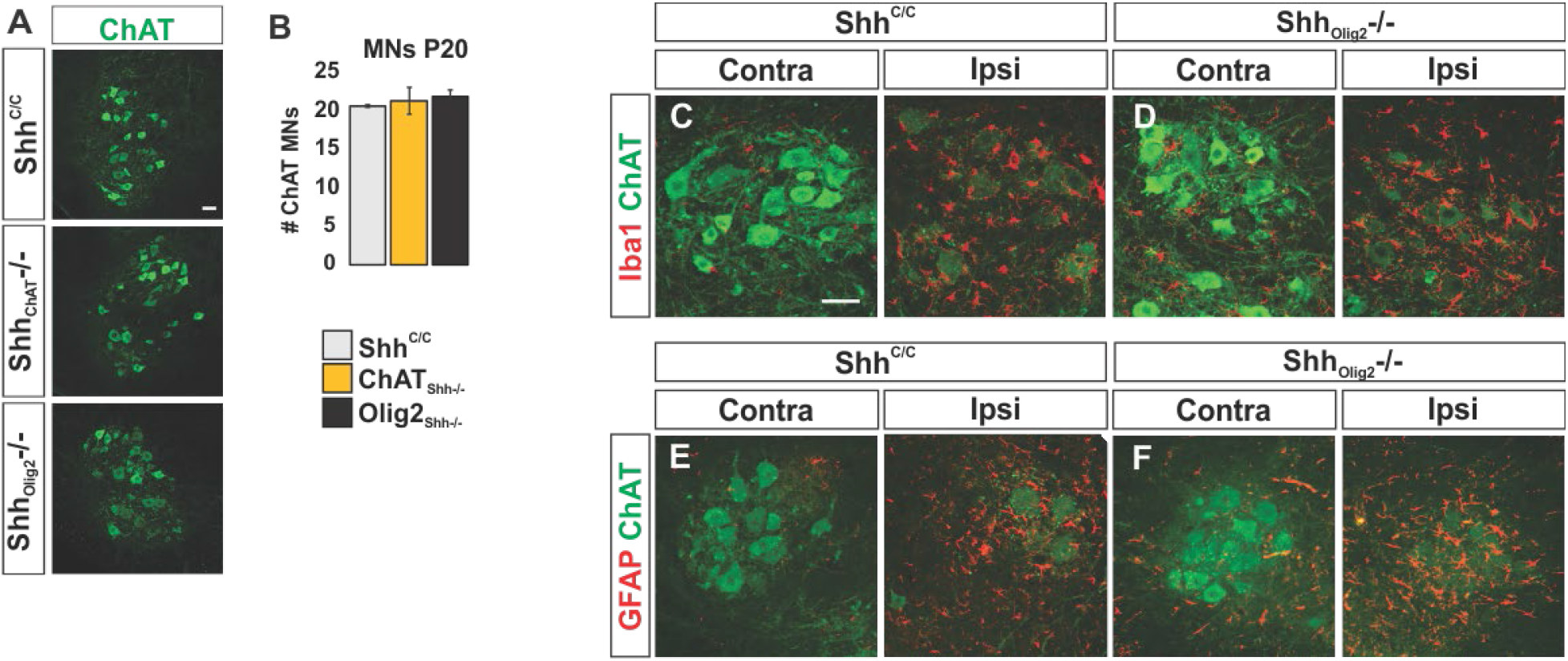
Microglia and astrocytes are activated in Olig2_Shh^−/−^_ mutants and controls in response to axotomy. (**A**) Lumbar motor neurons are organized and in normal numbers in ChAT_Shh^−/−^_ and Olig2_Shh^−/−^_ compared to control Shh^C/C^ (**B**) Quantification of MNs on L4-L5 lumbar spinal cord hemi sections. Shh^C/C^ (n = 4 mice), ChAT_Shh^−/−^_ (n = 4 mice), and Olig2_Shh^−/−^_ (n = 5 mice). Means ± SEM are shown. (**C and D**) Iba1+ microglia are recruited, reactive, and change morphology around injured ipsilateral MNs in Shh^C/C^ and Olig2_Shh^−/−^_. Scale bars, 50 μm. (**E and F**) GFAP+ astrocytes are recruited, reactive, and change morphology around injured ipsilateral MNs in Shh^C/C^ and Olig2_Shh^−/−^_.

